# Lipid composition determines thermotropic properties of meibum of animals and humans with Meibomian gland disorders

**DOI:** 10.1101/2025.08.25.672195

**Authors:** Igor A. Butovich, Jadwiga C. Wojtowicz, Amber Wilkerson, Seher Yuksel

## Abstract

Meibum – a lipid-rich secretion produced by holocrine Meibomian glands (MG) – plays a central role in maintaining ocular surface homeostasis. Previously, changes in MG lipidomes induced by inactivation of critical genes of meibogenesis, such as *Elovl3, Soat1, Awat2,* and others, were shown to cause MG-dysfunction- and dry eye-like signs in mice. Here, we describe the impact of the lipid composition of meibum on its physiological properties, specifically thermotropic/melting characteristics, using various wild type and mutant animals, and compare them with meibum of healthy humans and patients with abnormal meibum. Meibum samples were analyzed using liquid chromatography/mass spectrometry (LC/MS) and differential scanning microcalorimetry (DSC). We found that changes in the balance between major lipid classes in meibum – wax esters, cholesteryl esters, triacylglycerols and free cholesterol – cause detrimental changes in its properties, leading to the loss of cohesiveness, abnormal expressibility from MG, a decline in its protective functions, and, ultimately, abnormal phenotypes of the eyes and adnexa. We conclude that tested animal species can be valuable models for studying human diseases. A combination of LC/MS and DSC can be a powerful diagnostic tool and may help to determine the molecular reasons of various pathologies in human subjects and animals.

## 1. INTRODUCTION

Sebum and meibum are two types of lipid-rich secretions that are produced by related, but not identical types of exocrine/holocrine sebaceous and Meibomian glands (SG and MG, correspondingly). Sebum creates a protective hydrophobic layer that covers the skin, human hair, and animal fur^1^, while meibum^2^ is one of the main components of the preocular tear film^3^. The main functions of sebum and meibum are similar: sebum is a lubricant that protects the skin from desiccation and exogenous pathogenes, while meibum is performing similar functions in the eye. To work, sebum and meibum need to be expressed from their corresponding glands as a first step. This mechanism involves the (mostly) passive transport of the secretions through a system of ducts before being released onto the skin or the ocular surface. The ability of the secretions to passively travel through the ducts heavily depends on their composition: secretions that are too thick may severely reduce the flow and cause stagnation of sebum and meibum in the ducts and glands, while secretions that are abnormally fluid will cause the lipid overflow – conditions that are detrimental to both the ocular surface and the skin homeostasis.

Indeed, deviations in the chemical composition and/or quantity of biosynthesized meibum from the norm due to pathologies known as hypo- and hypersecretory MG dysfunctions (MGD), have been reported to trigger the onset, or exacerbate the consequences of, dry eye (DE) ^4–6^. Similar effects are expected to play a role in the release of abnormal sebum from free and hair-associated SG.

Previously, targeted changes in MG lipidomes induced by inactivating mutations in critical genes of meibogenesis^7^, such as *Elovl3, Soat1, Awat2, Cyp4f39*, *Sdr16c5/Sdr16c6, Scd1, Far1/2,* and others, were shown to almost universally cause MGD- and DE-like signs in mice^8–16^. In this paper, we describe the impact of differences in the lipid composition of mouse, canine, and rabbit meibum on its biophysical/physiological properties, specifically thermotropic/melting characteristics, and compare them with meibum and sebum of healthy human volunteers and, patients with abnormal meibum.

## 2. RESULTS

### 2.1. Untargeted, unsupervised LC/MS and multivariate statistical analyses of human and animal meibum

Chemical composition of meibum varied significantly between the species and genotypes. Results of the initial evaluation of the human and animal Meibomian lipidomes were reported earlier^17^. However, for the purpose of this study, we re-analyzed human and animal meibum using liquid chromatography/high resolution time-of-flight mass spectrometry (LC/MS) and compared the samples using unbiased, untargeted, unsupervised Principal Component Analysis (PCA) and supervised Partial Least Squares Discriminant Analysis (PLS-DA) approaches included in Progenesis QI and EZinfo software packages (all from Waters Corp., Milford, MA, USA). The data are shown in Figure 1A as a Loadings Bi-Plot. Note a rather tight intra-group clustering of the samples of the same origin (shown as circles of different colors), and clear inter-group separation of the samples. The black dots represent the lipid analytes, whose proximity to specific groups of samples depends on their associations with specific species or genotypes. The chemical composition of meibum varied significantly between the species and genotypes, but a close proximity of human and canine groups on the plot is indicative of their close biochemical similarities. Wild type (WT) mice also demonstrated rather tight grouping of the samples of the same type, and were the second closest group to human samples. Meibum samples of *Sdr16c5^−/−^/Sdr16c6^−/−^*double knockout (*DKO*) mice were the most distant from both WT mouse and human meibum. The rabbits are not included in the PCA/PLS-DA analyses because of their fundamental differences from other species: Rabbit meibum is based primarily on dihydrolanosteryl esters (DiHLE) as major nonpolar lipid components, and much smaller amounts of cholesteryl esters (CE) and wax esters (WE), which are major Meibomian lipids in other studied animals^17^.

**Figure 1.**
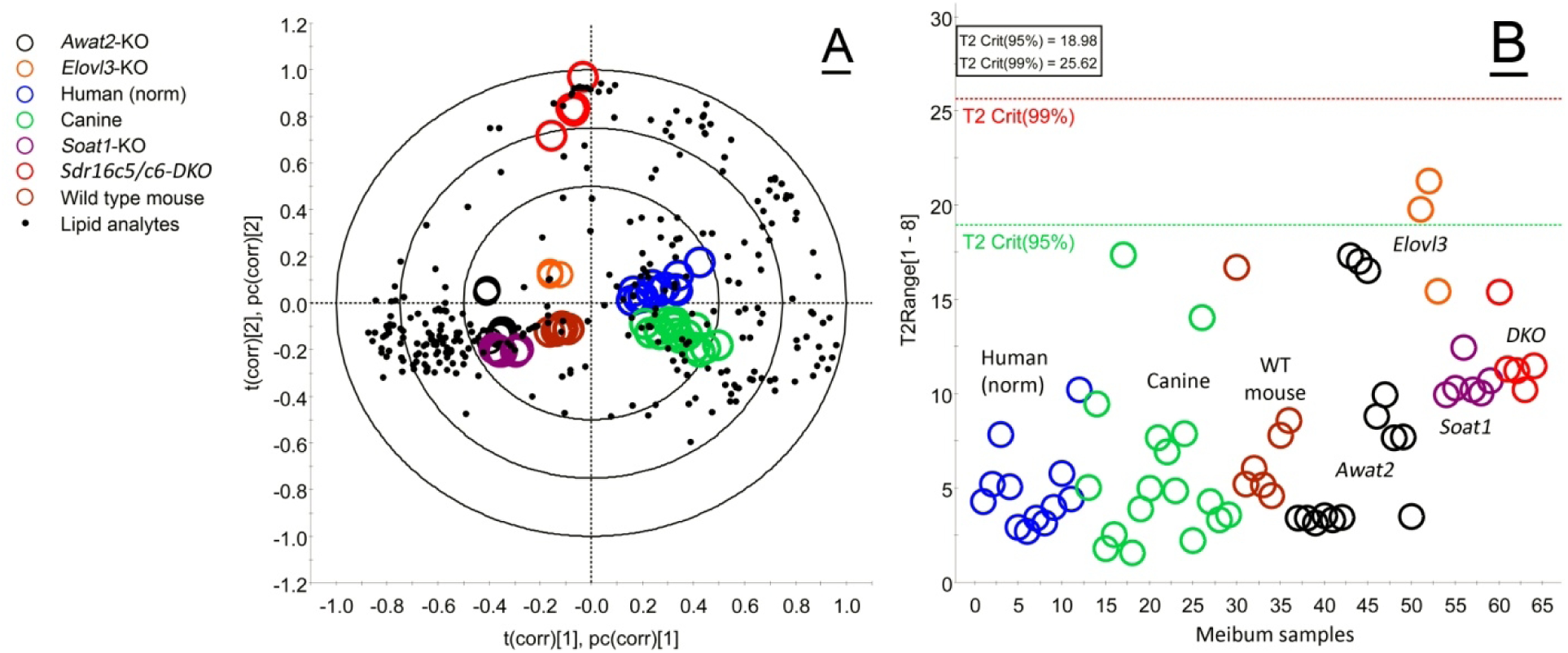
Multivariate statistical analysis of meibum samples obtained from different human subjects and animal species highlighted their similarities and differences. *<u>Panel A.</u>* Partial Least Squares Discriminant Analysis (PLS-DA) of study samples (shown as colored circles) demonstrated their clear separation based on the origin, and rather tight intra-group clustering. The lipid analytes are shown as black dots. *<u>Panel B.</u>* The Hotelling’s T^2^ plot was used to detect critical outliers in the study groups that would exceed the T^2^ Crit(99%) threshold, but none was detected.

Importantly, the Loadings Bi-Plot also revealed the type of lipids that were associated with specific groups of samples (discussed below), while the Hotelling’s T^2^ Range Plot demonstrated that there were no significant outliers in the datasets: all specimens were well below the T^2^ Crit(99%) threshold, with only two samples barely exceeding the T^2^ Crit(95%) threshold (Figure 1B).

### 2.2. Mouse meibum

#### 2.2.1 Characterization of the chemical composition of wild type and mutant mouse meibum

Then, more targeted comparative liquid chromatography/mass spectrometry (LC/MS) analyses of wild type mouse and normal human meibum were performed, to ensure consistency of the results with previously published data^7, 8, 10, 12, 17–19^.

First, the effects of the mutations on mouse Meibomian lipidomes were studied (Figure 2). The wild type (*WT*) meibum of C57BL/6J mice was used as a baseline for comparing the effects of mutations on the composition and properties of meibum of genetically modified mice.

**Figure 2.**
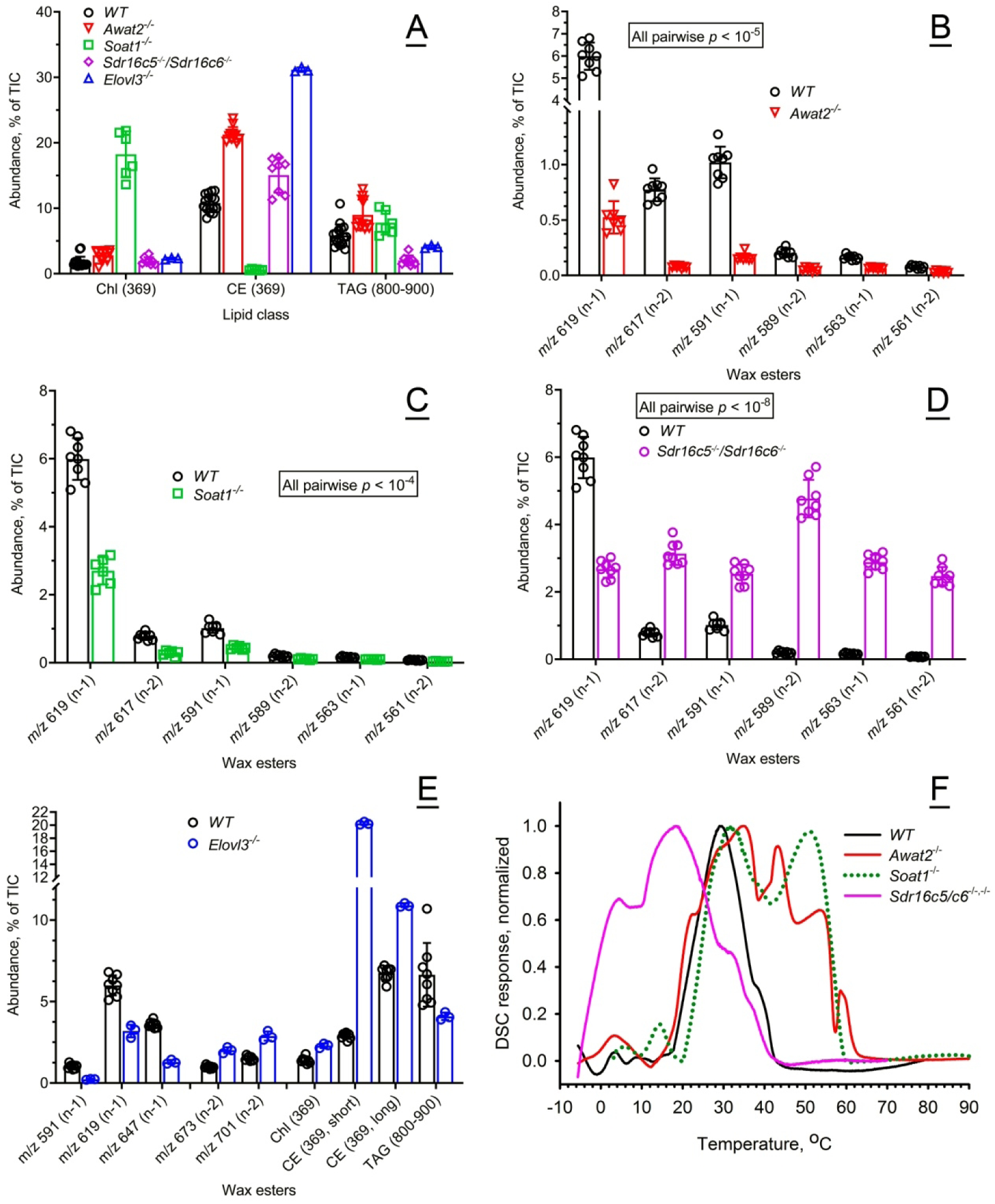
Compositional and functional characterization of mouse meibum from wild type (*WT*) and mutant mice. *<u>Panel A.</u>* Relative balance of free cholesterol (Chl), cholesteryl esters (CE), and triglycerides (TAG) in meibum of wild type (*WT*) and mutant mice. The balances were calculated by calculating the peak areas of the extracted ion chromatograms of the lipids as shown in Figure 1. The data is shown as percentage points of total lipids. *<u>Panel B.</u>* Relative balance of major wax esters detected in *WT* and *Awat2^−/−^* mouse meibum. *<u>Panel C.</u>* Relative balance of major wax esters detected in *WT* and *Soat1^−/−^* mouse meibum. *<u>Panel D.</u>* Relative balance of major wax esters detected in *WT* and *Sdr16c5^−/−^/Sdr16c6^−/−^* mouse meibum. *<u>Panel E.</u>* Relative balance of major wax esters, Chl, short- and long-chain CE, and TAG detected in *WT* and *Elovl3^−/−^* mouse meibum. *<u>Panel F.</u>* Normalized thermograms of *WT* and mutant mouse meibum demonstrated critical and selective impact of the mutations on the melting characteristics of the secretions. Representative samples are shown. The averaged data for tested samples can be found in Table 1.

**Table 1.** Thermotropic properties of human and animal meibum.

| *Animals and human subjects* | *Ocular and skin abnormalities* | *Major features of MG lipidomes* | $T_m$, °C<br>(Mean ± SD)* | $T_{1/2}$, °C<br>(Mean ± SD)* |
| --- | --- | --- | --- | --- |
| C57Bl6/J mice | None | Similar to human | 28 ± 1 | 19 ± 3 |
| *Awat2*-knockout mice | Severe ocular and Meibomian gland phenotype; solid meibum | WE↓, free Chl↑ and CE↑ | Multiple melting peaks between 20 and 70 °C | Extremely wide, >40 °C |
| *Soat1*-knockout mice | Severe ocular and Meibomian gland phenotype; solid meibum | Lacks CE, free Chl↑↑↑ | Biphasic melting curve;<br>30 ± 2 (1<sup>st</sup>)<br>50 ± 1 (2<sup>nd</sup>) | Each peak > 20 °C |
| *Elovl3*-knockout mice | Severe ocular and Meibomian gland phenotype; liquefied meibum | Abnormal WE and CE elongation and unsaturation patterns; ELC WE↓ | Very low, < 0 °C | n/d |
| *Sdr16c5/Sdr16c6* double knockout mice | Severe ocular and Meibomian gland phenotype; liquefied meibum | Transformed into a sebum-like lipidome; TAG↑ and short WE↑ | 19 ± 1 | 30 ± 5 |
| Healthy canines | None | Similar to human | 29 ± 1 | 13 ± 2 |
| White New Zealand rabbits | Abnormally stable tear film | Based primarily on DiHL, DiHLE, DiAD, SE (not CE), and OAHFA | 33 ± 1 | 8 ± 1 |
| Normal human subjects (HS) | None | Based primarily on ELC saturated and mono-unsaturated WE, CE, free Chl, TAG, OAHFA, Chl-OAHFA, and DiAD | 29 ± 1 | 10 ± 1 |
| Abnormal HS #1 | Dry eye, skin and solidified meibum | Saturated WE↑↑↑ | 39 ± 1 | 12 ± 1 |
| Abnormal HS #2 | Lard-like, solidified meibum; abnormal tear film lipid layer with lipid rafts | TAG↑, short WE↑, ELC WE↓ | Biphasic melting curve;<br>21 ± 2 (1<sup>st</sup>)<br>57 ± 2 (2<sup>nd</sup>) | ~17 (1<sup>st</sup>)<br>~15 (2<sup>nd</sup>) |

\* The values were rounded up or down to the nearest integral number; averaged numbers for multiple samples are shown

Inactivating mutations in *Elovl3*, *Soat1, Awat2,* and *Sdr16c5/Sdr16c6* genes in the mice on the same background led to fundamental shifts in their Meibomian lipidomes, affecting the ratios between individual lipids within their respective groups, and balance between different lipid groups and classes. As meibum is predominantly based on nonpolar lipids, such as WE, CE, free cholesterol (Chl), and triacylglycerols (TAG), these lipids were profiled in the study samples as shown in Figures 2A-2E. The levels of Chl were severely upregulated by inactivation of *Soat1*, while inactivation of other genes had only minimal impact on the compound. On the other hand, CE as a group were almost nonexistent *Soat1^−/−^* mice, considerably upregulated in *Awat2^−/−^* mice, but very close in *WT* and *Sdr16c5^−/−^/Sdr16c6^−/−^ DKO* mice. The levels of TAG, which are the main sources of carbon and energy for lipogenesis in cells, were almost unaffected in *Awat2^−/−^*and *Soat1^−/−^*mice, but somewhat suppressed in other mutants.

Inactivation of *Awat2* in mice has been previously reported to lead to a decline^26^, or an almost complete suppression^10, 20^, of the biosynthesis of Meibomian WE in mouse tarsal plates. This observation was reconfirmed for expressed *Awat2^−/−^* meibum in our current experiments (Figure 2B) – a multifold decrease in the levels of mono- and di-unsaturated Meibomian WE (MUWE and DUWE, correspondingly) was observed, which, together with a concomitant increase in Chl and CE, caused a fundamental change in the MG lipidome. Therefore, the remaining major components of the *Awat2^−/−^* meibum were Chl, CE, and related compounds.

Aside from an almost complete suppression of formation of CE and upregulation of Chl, inactivation of *Soat1* lowered the expression levels of tested WE, making Chl the major lipid in meibum (Figures 2A and 2C). While the overall ratios of Chl, CE, and TAG in meibum of *DKO* mice underwent only modest alterations, the pool of their nonpolar lipids became highly enriched with sebaceous-type lipids, such as diunsaturated and shorter-chain WE (Figure 2D). Note that the *total* amount of biosynthesized meibum in the *DKO* tarsal plates was ∼3× of that in *WT* mice, so the *total* fraction of Meibomian-type WE in meibum did not change much, while the increase in WE was caused by accumulation of sebaceous-type WE^19, 21^.

Inactivation of *Elovl3* changed the balance between MUWE and DUWE toward the latter, and led to a considerable increase in the abundance of shorter-chain CE, with fatty acid (FA) chains ≤ C_20_), a ∼ 30% increase in their longer chain counterparts (FA > C_20_), and a ∼40% decline in TAG (Figure 2E).

#### 2.2.2. Effects of the mutations on thermotropic properties of wild type and mutant mouse meibum

For the reader’s convenience, the results of differential scanning microcalorimetry (DSC) experiments with mouse meibum are summarized in Table 1 and Figure 2F. Close similarities between human and *WT* mouse meibum were expected to result in close thermotropic properties of both secretions, with, possibly, slightly lower transition temperature (**T_m_**) of the mouse meibum. Indeed, mouse meibum produced a well-defined, rather symmetrical thermogram, and had a major apparent **T_m_** between 26 and 28 °C, which was ∼2 °C lower than the human one, and a somewhat broader major melting range with a **T_1/2_** (the width of the transition at half-height of the melting peak) of 19 ± 3 °C. The broadening of the T_1/2_ was caused mainly by the ascending part of the melting curve which showed a first sign of a phase transition between −5 °C and 0 °C. Interestingly, its upward section had a few faint shoulders which were either much less pronounced, or non-existent, in human samples, and might indicate either heterogeneity of the lipid mixture, or minute phase transitions in its structure. Another distinctive feature of mouse meibum was a shoulder, or in some cases, a minor transition peak, with a T_m_ of around 38 °C to 40 °C. However, the downward section of the curve had almost the same shape and slope as the human one. The most distinctive feature of the *WT* mouse meibum was a secondary, though minor, transition peak, or a shoulder in some samples, with a T_m_ of around 40 °C. *WT* mouse meibum became completely liquefied at temperatures above 45 °C. The significance of these features for the ocular surface physiology will be discussed later in the paper.

Changes in the chemical composition of meibum caused by the mutations radically changed its melting properties (Figure 2F) and, hence, expressibility from MG orifices.

The *Awat2* mutation had an unexpectedly complex effect on the thermotropic properties of the mutant meibum with multiple transitions/calorimetric domains being observed. Aside from a low-temperature transition peak with T_m_ of ∼3 °C, multiple transitions between T_m_ of ∼35 °C and ∼58 °C were observed with an overall T_1/2_ of more than 35 °C – the widest among all animal models tested in this study.

The thermotropic properties of *Soat1^−/−^* meibum were also fundamentally different from those of *WT* mice. The melting curves were of bimodal nature, though three apparent transition temperatures were observed: 14 ± 1 °C (minor), 32 ± 1 °C (major), and 51 ± 1 °C (major). The estimate value of T_1/2_ was also wide at ∼30 °C.

Commensurate with the shorter-chain, and less saturated, nature of Meibomian lipids, their melting started at below −5 °C, with a major apparent T_m_ of ∼18.2 °C being about 10 °C lower than the T_m_ of *WT* meibum (Figure 2F). Interestingly, a shoulder with a T_m_ of around 3 °C was visible and close to the similar low temperature transition peaks in *WT* and *Soat1^−/−^* mice. The T_1/2_ of *DKO* meibum ( 28.5 ± 1.5 °C) was considerably wider than the T_1/2_ of *WT* mice (19 ± 3 °C).

### 2.3. Canine meibum

Canine MG secretions are studied to a much lesser extent than the human and mouse ones, though being house pets, their ocular conditions are closely monitored and treated by veterinarians^22–26^. Results of initial characterization of the chemical composition of canine meibum by means of ion trap mass spectrometry (MS^n^) had been described earlier^17, 27^. The secretions were re-evaluated in the current study using a more accurate LC/MS (Figure 3). Total ion chromatograms of both secretions are shown in Figure 3A, while their observation mass spectra – in Figure 3B. The major canine WE were present with a very distinctive pattern – *m/z* 605#, *m/z* 619∃, *m/z* 633# (major peak), *m/z* 647∃, *m/z* 661# – while human WE showed an opposite trend – *m/z* 605∃, *m/z* 619#, *m/z* 633∃, *m/z* 647# (major peak), *m/z* 661∃. The individual components in each type of samples were compared using extracted ion chromatograms (EIC) as shown in Figure 3C, and their identities were established on the basis of their *m/z* values, chromatographic retention times, and MS/MS fragmentation patterns. Canine and human WE have similar saturation/desaturation ratios being based primarily on C_16:0_, C_18:0_, C_16:1_, and C_18:1_ fatty acids and saturated fatty alcohols. Also, canine and human CE and free Chl were present in comparable (but not identical) amounts and ratios^17, 27^.

**Figure 3.**
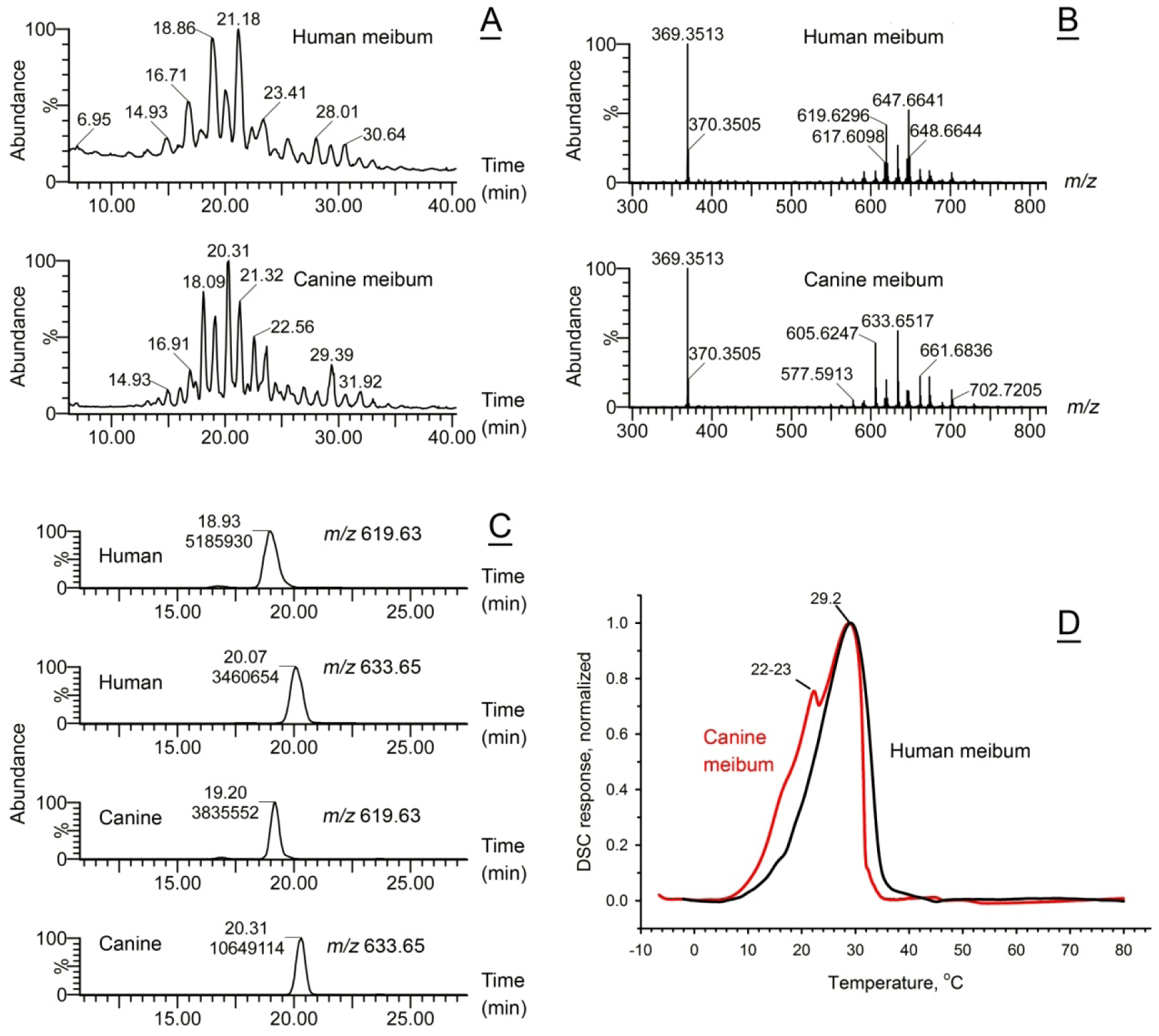
Compositional and functional characterization of canine meibum demonstrated its close similarity with normal human meibum. *<u>Panel A.</u>* Total APCI-PIM ion chromatograms of representative human (upper trace) and canine (lower trace) meibum samples obtained as shown in Figure 1, Panel A. *<u>Panel B.</u>* Observation APCI-PIM mass spectra of the human and the canine samples shown in Panel A. *<u>Panel C.</u>* Extracted ion chromatograms of two major wax esters in human (two upper traces) and canine (two lower traces). *<u>Panel D.</u>* Normalized thermograms of human and canine meibum samples demonstrated close similarities between the melting characteristics of the secretions. A minor peak, or a shoulder in some canine samples, was responsible for their slightly wider melting peaks. Representative samples are shown. The averaged data for tested samples can be found in Table 1.

Surprisingly, despite the reversed WE expression patterns and other differences, the major apparent transition temperature of canine meibum – about 29 ± 1.5 °C – was extremely close to that of a representative human sample, while the T_1/2_ value was ∼12 ± 1 °C – again, almost the same as for normal human meibum (Figure 3D). A minor shoulder at around 22-23 °C, detected in some canine samples, was a feature that differentiated them from human meibum, though its low intensity and rather low T_m_ (which is well below physiological levels) seems to be less relevant to the ocular surface physiology.

### 2.4. Rabbit meibum

Rabbits are popular laboratory animals that are frequently used in biomedical experiments and pharmaceutical trials due to the anatomical similarities of their ocular structures with human eye^26, 28, 29^ ^30^. However, the lipidomic profile of rabbit meibum is fundamentally different from that of humans, canines, and WT mice^17^. Rabbit meibum is composed primarily of a rather complex mixture of dihydrolanosteryl (DiHL) esters (DiHLE), steryl esters (SE) (Figures 4A-4C), and some other compounds of polar and nonpolar nature. Because of its complexity, comprehensive characterization of rabbit meibum has not been performed yet and is left for future studies. Interestingly, most of the major lipids in rabbit meibum are not found in humans; however, despite being fundamentally different from human meibum, the T_m_ of the rabbit secretion is only 3 to 4 degrees higher than the human one (Figure 4D). The T_1/2_ value of the rabbit meibum melting peak was between 7 and 8 °C, a few degrees less than the human and canine secretions, implying higher cooperativity of melting and sharper responses to the changes in temperature.

**Figure 4.**
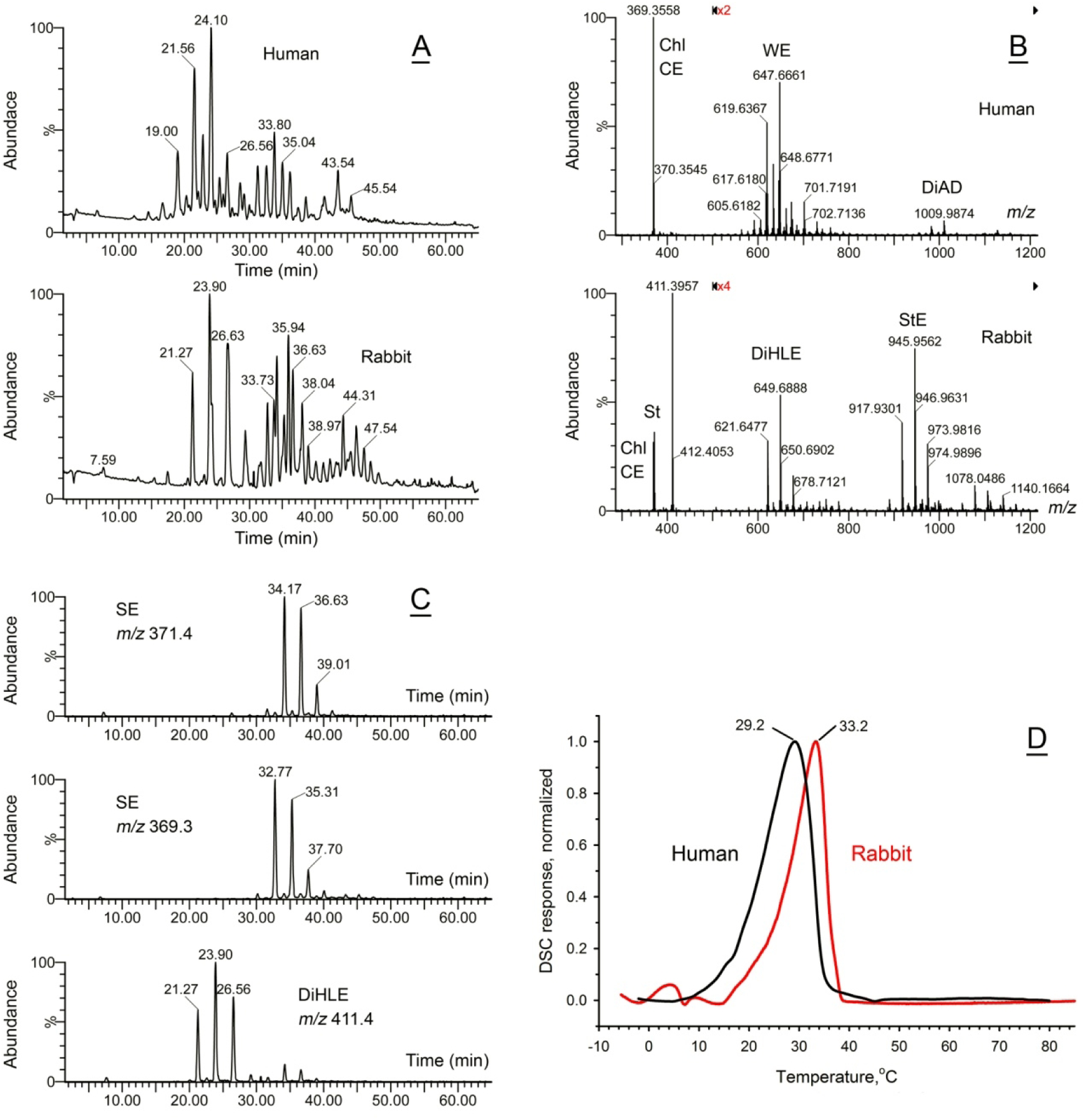
Compositional and functional characterization of rabbit meibum. *<u>Panel A.</u>* Total APCI-PIM ion chromatograms of representative human (upper trace) and rabbit (lower trace) meibum samples obtained as shown in Figure 1, Panel A. *<u>Panel B.</u>* Observation APCI-PIM mass spectra of the human and the rabbit samples shown in Panel A illustrate their fundamental biochemical differences. *<u>Panel C.</u>* Extracted ion chromatograms of typical steryl esters in rabbit meibum. *<u>Panel D.</u>* Normalized thermograms of human and rabbit meibum samples demonstrated a surprising similarity of the shapes of their thermotropic peaks, albeit a somewhat higher melting temperature of the rabbit meibum. Representative samples are shown. The averaged data for tested samples can be found in Table 1.

### 2.5. Normal and abnormal human meibum

Then, meibum from human subjects was studied using untargeted and targeted LC/MS experiments. Truly quantitative analysis of all, or even major only, Meibomian lipids is yet to be completed due to the lack of necessary lipid standards for a wide variety of Meibomian lipids. Therefore, for the purpose of this study, the apparent LC/MS abundances of major detected analytes were used, compared, and discussed instead.

#### 2.5.1. Normal human meibum

Normal human meibum has been qualitatively and semi-quantitatively characterized on the levels of lipid classes and individual lipid species by LC/MS and gas chromatography/MS (GC/MS) in previous studies performed by our laboratory and by independent teams. It has been firmly established that the major components of meibum are numerous WE and CE, free Chl, free fatty acids (FFA), TAG, (*O*)-acylated ω-hydroxy fatty acids (OAHFA), cholesteryl esters of OAHFA (Chl-OAHFA), diacylated α,ω-diols (DiAD), small amounts of phospholipids and sphingomyelins, and other minor compounds of lipid nature^31–36^.

First, human meibum samples were collected from normal, non-MG dysfunction (MGD), non-dry eye (DE) volunteers and individually analyzed using LC/MS. Untargeted analyses of normal human meibum samples revealed their rather tight grouping with no clear differentiating factors – note large SD values even for the most influential components of the secretions, and lack of outliers, which correlated well with their highly similar DSC thermograms (Figures 5A-5D).

**Figure 5.**
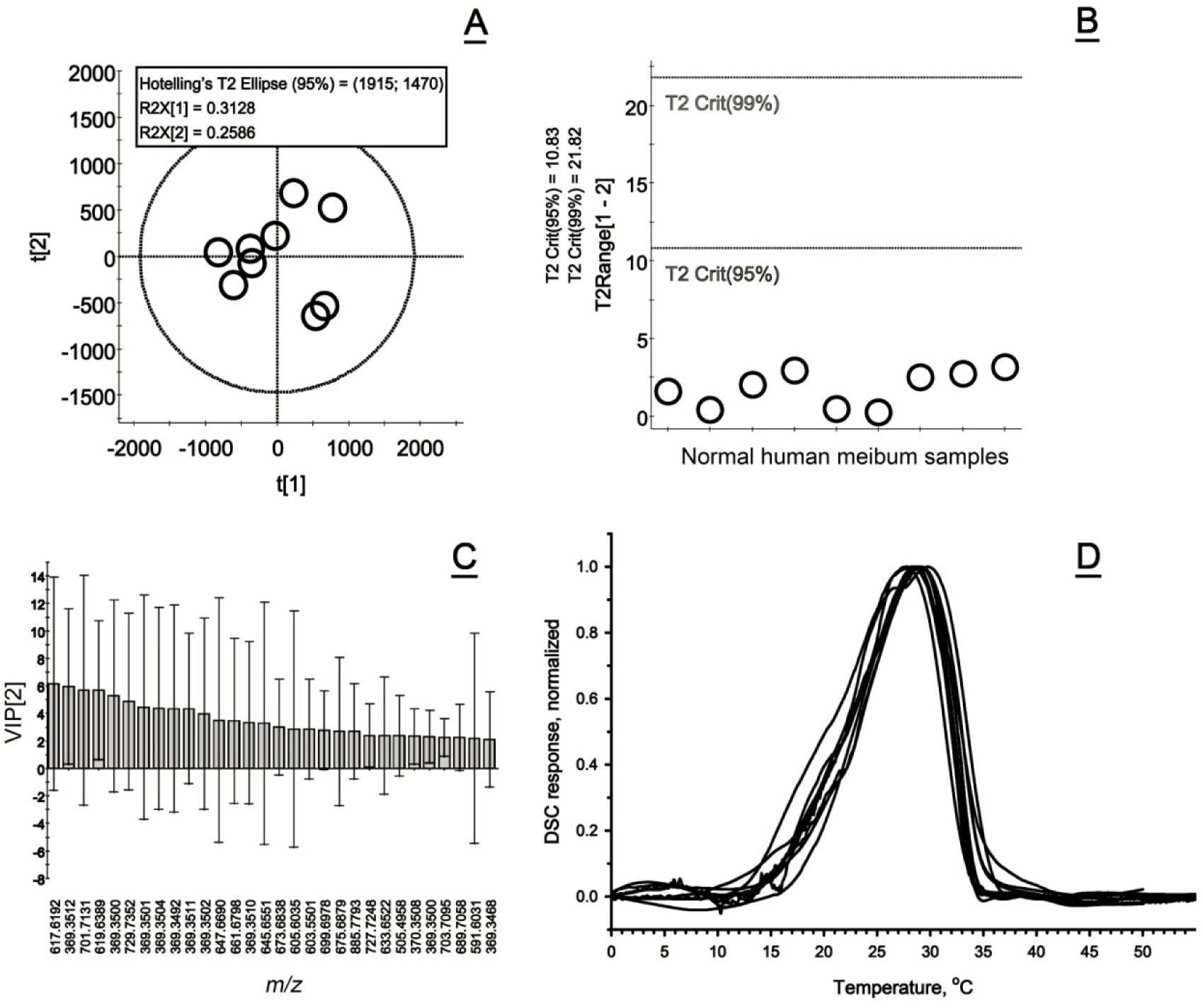
Compositional and functional characterization of normal human meibum. *<u>Panel A.</u>* A Principal Component Analysis Scores plot of normal human meibum samples analyzed as shown in Figure 1. *<u>Panel B.</u>* No outliers were detected using the Hotelling’s T^2^ plot as none of the samples approached the T^2^ Crit(95%) threshold when analyzing permutated data using PLS-DA template. *<u>Panel C.</u>* The Variable Importance plot shows 30 most influential lipid analytes (mostly, wax esters, cholesteryl esters, and triacylglycerols) with the highest impact on separation of the samples. However, the differences between the expression levels of these lipids were minimal and statistically insignificant. *<u>Panel D.</u>* Normalized thermograms of 11 normal human meibum samples collected from different subjects demonstrated similar melting behavior with close T_m_ and T_1/2_ values (Table 2).

**Table 2.** Interdonor variability of the thermotropic properties of normal human meibum.

| *Human subject* | *Gender*<br>*(Age, years)* | *Meibum*<br>*$T_m, ^\circ\text{C}, *$* | *Meibum*<br>*$T_{1/2}, ^\circ\text{C}^*$* |
| --- | --- | --- | --- |
| 1 | M (41) | 29.2 | 10.9 |
| 2 | M (56) | 28.6 | 9.9 |
| 3 | F (48) | 30.9 | 10.2 |
| 4 | M (25) | 29.1 | 9.3 |
| 5 | F (41) | 27.8 | 11.2 |
| 6 | not<br>recorded | 29.6 | 10.7 |
| 7 | F (42) | 28.5 | 9.3 |
| 8 | F (27) | 28.3 | 9.9 |
| 9 | F (28) | 27.7 | 11.0 |
| 10 | F (40) | 28.6 | 9.62 |
| 11 | F (60) | 28.1 | 10.6 |
| Average (mean ± stand. dev.) | 41 ± 11 | 28.8 ± 0.9 | 10.2 ± 0.7 |

\* The values were rounded up or down to the nearest integral number

Targeted analyses of specific lipids and lipid classes were conducted using EIC of specific lipids as shown in Figure 6 for Chl, CE, some of major Meibomian WE, and a major TAG triolein. Two different LC methods were used – a gradient LC on a C18 BEH reverse phase column (C18-RP LC) and isocratic LC on a reverse phase C8 BEH column (C8-RP LC). The first approach was used for detailed analyses of Meibomian lipidomes, while the second – for lipid quantitation (Figures 6A and 6B)^19^. All LC/MS experiments were conducted in the atmospheric pressure chemical ionization positive ion mode (APCI PIM). A representative observation mass spectrum of normal human meibum sample is shown in Figure 6C, which provided the data for analyzing the major components of meibum – WE, CE, Chl, DiAD and other lipids as depicted in Figure 6D.

**Figure 6.**
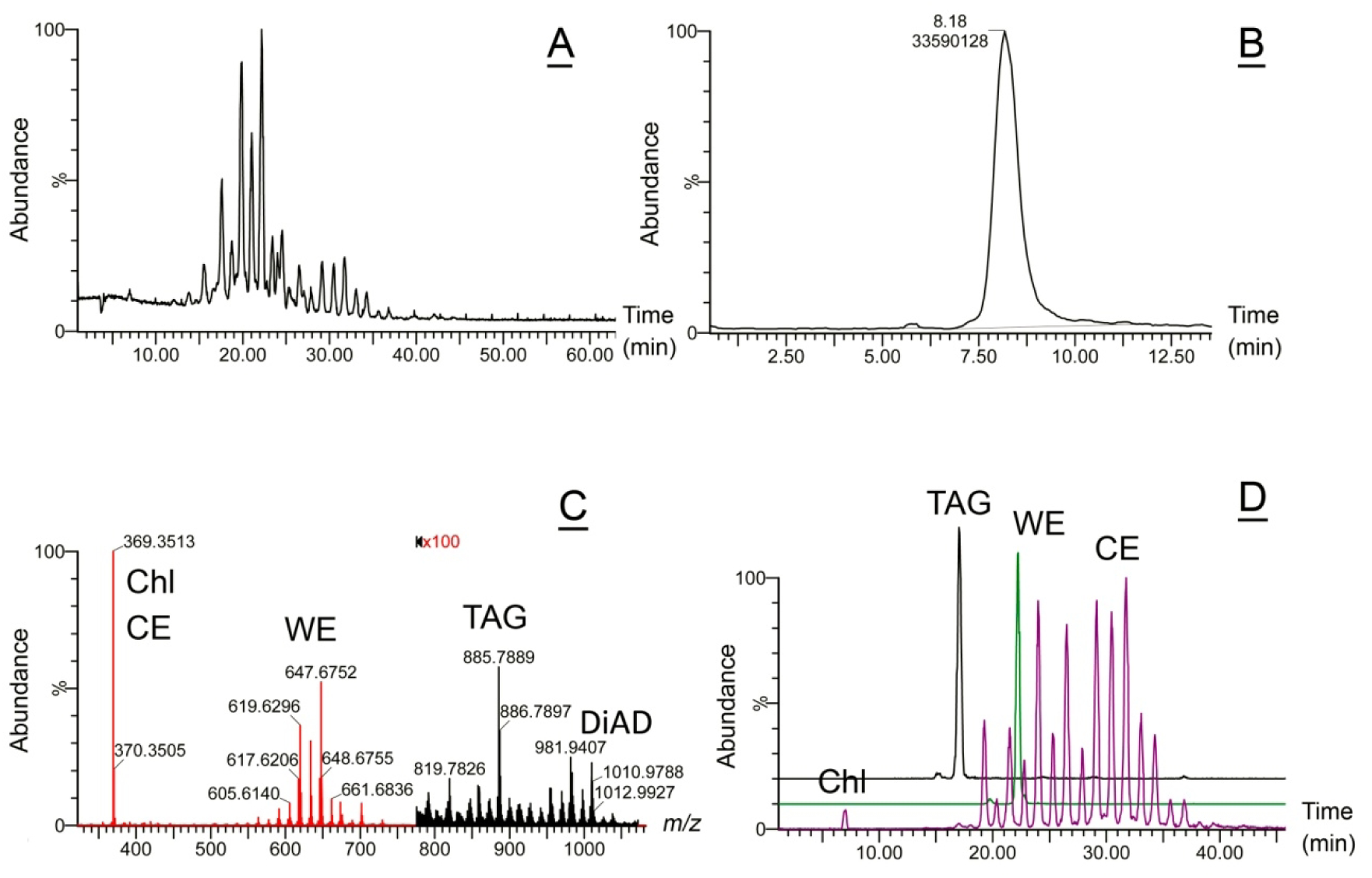
Targeted lipidomic analysis of normal human meibum using liquid chromatography/mass spectrometry (LC/MS) reveals major types of Meibomian lipids. *<u>Panel A.</u>* A total ion chromatogram of a representative sample of human meibum. Conditions of the LC/MS analysis were as described before^19^: a C18 BEH LC column; gradient elution with iso-propanol/acetonitrile/aqueous ammonium formate solvent mixture; atmospheric pressure chemical ionization mass spectrometry in positive ion mode (APCI PIM). Meibomian lipids eluted as multiple LC/MS peaks with retention times between 5 and 55 min. *<u>Panel B.</u>* Meibomian lipid quantitation using isocratic LC/MS. Conditions of the LC/MS analysis: a total ion chromatogram of the same sample as in Panel A was obtained using iso-propanol/acetonitrile/aqueous ammonium formate solvent mixture and a C8 BEH LC column^19^ and APCI-PIM. Meibomian lipids eluted as a major LC/MS peak with a retention time of 8 ± 1 min, with a much smaller peak at around 6 ± 1 min. Integration of the major peak provided an estimate of the lipid content in the sample in relative units. *<u>Panel C.</u>* An observation APCI-PIM mass spectrum of the human sample shown in Panel B. Note that the abundance of triacylglycerols (TAG) and diacylated α,ω-diols (DiAD) in the sample was very low which necessitated 100× magnification of that part of the spectrum, to show their signals. *<u>Panel D.</u>* Extracted ion chromatograms (EIC) of triolein (TAG, black trace), one of major Meibomian wax esters C_42_H_82_O_2_ (WE, green trace), free cholesterol and cholesteryl esters (Chl and CE, both purple trace). Conditions of the LC/MS analysis were the same as in Panel A.

In total, eleven normal meibum samples were analyzed by DSC (Table 2). Due to the sensitivity limits of the instrumentation, only samples that exceeded 0.2 mg were selected for the experiments. Normal meibum produced thermograms that had a main average T_m_ at around 28.8 ± 0.9 °C and a T_1/2_ of 10.2 ± 0.7°C (Figure 5D). Their melting started at around 12 ± 2 °C. Compared with pure lipids^37^, meibum thermograms showed rather broad melting peaks with a slightly shallower ascending and steeper descending portions. As evidenced by the thermograms, meibum became solidified at <10 °C, and completely liquefied at a physiological body temperature of 37 ± 2 °C, being in a partially melted state at a physiological cornea temperature of 32 ± 2 °C ^38–42^. We consider this melting behavior in the ∼30 °C to ∼37 °C temperature range to be absolutely for proper expression of meibum from MG orifices and its protective properties (see the Discussion section for details).

#### 2.5.2. Abnormal human meibum

Abnormal meibum samples collected from human Subjects **<u>1</u>** and **<u>2</u>** were studied using the same LC/MS and DSC approaches as described above. Initially, both types of abnormal samples were compared with normal human meibum using unbiased, untargeted PLS-DA approaches. The three types of samples formed tight, but clearly separated groups, of which none of the samples was an outlier – all samples fell below the T^2^ Crit(95%) threshold (Figures 7A and 7B).

**Figure 7.**
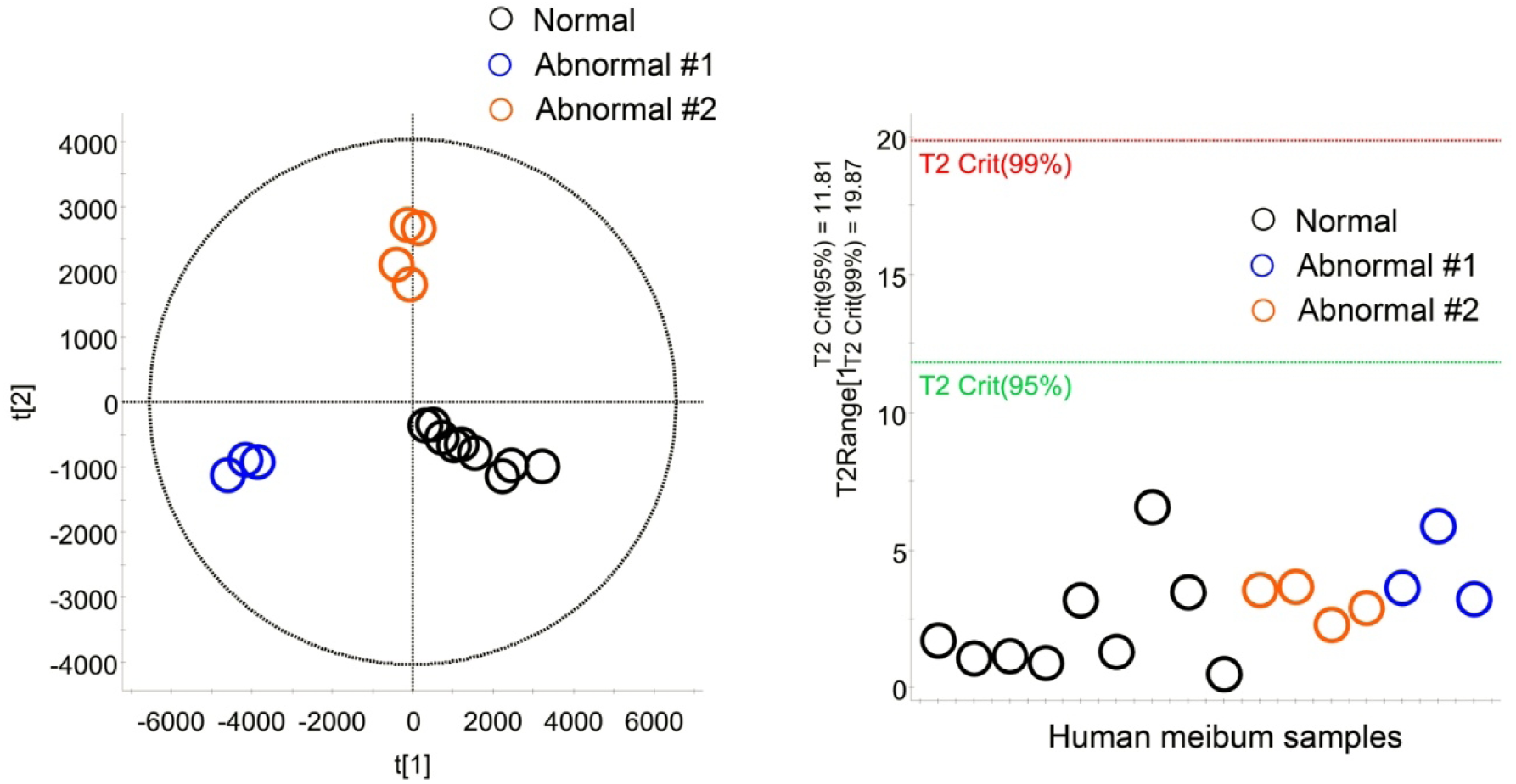
Untargeted, unbiased comparison of normal and abnormal human meibum. *<u>Panel A.</u>* A Principal Component Analysis Scores plot of normal (N) and abnormal (#1 and #2) human meibum samples analyzed as shown in Figure 1. *<u>Panel B.</u>* No outliers were detected using the Hotelling’s T^2^ plot as none of the samples approached the T^2^ Crit(95%) threshold.

*Subject #1.* The Meibomian lipid profiles and ocular surface features of Subject **<u>1</u>** have already been partially characterized in our earlier publication^43^. Subject **<u>1</u>**’s meibum samples were collected on three different occasions a year apart from each other, and analyzed in this study alongside other human and animal specimens. The experiments produced the same result as before (Figures 8A and 8B): a severe imbalance between WE, CE, Chl, and TAG. Specifically, the lipid profile of Subject **<u>1’s</u>**meibum samples resembled that of human sebum being highly enriched with shorter-chain, and less saturated, WE compared to normal meibum [Figure 8C and (<u>5</u>)], among other changes.

**Figure 8.**
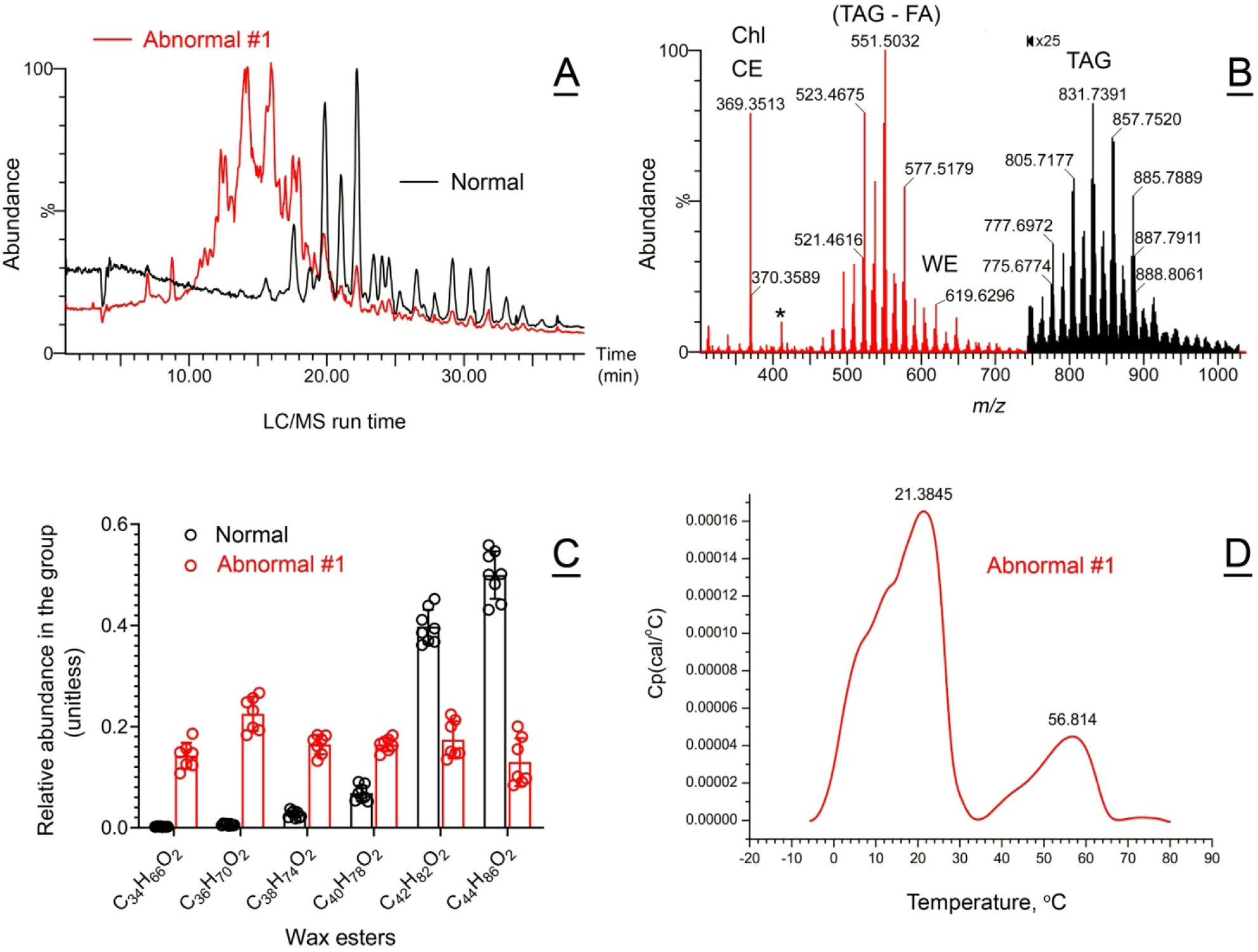
Compositional and functional characterization of abnormal human meibum #1. *<u>Panel A.</u>* Total APCI-PIM ion chromatograms of representative abnormal #1 (orange trace) and normal (black trace) meibum samples obtained as shown in Figure 1, Panel A. The vast majority of the signals detected in abnormal sample #1 came from a pool of faster eluting, less hydrophobic analytes. *<u>Panel B.</u>* Observation APCI-PIM mass spectrum of abnormal human sample #1. Note that the TAG region of the spectrum was magnified ×25 times for clarity. *<u>Panel C.</u>* Shorter-chain, sebum-like wax esters were highly upregulated in abnormal sample #1, at the expense of longer-chain, Meibomian-type wax esters. *<u>Panel D.</u>* Abnormal meibum melted in a bimodal manner with two clearly different T_m_ of 21 and 57 °C.

DSC experiments with Subject **<u>1’s</u>** meibum samples produced a clearly abnormal biphasic/bimodal pattern with two major apparent melting peaks with T_m_ of 21 ± 1 °C and 57 ± 2 °C, and several shoulders (Figure 8D). Notably, melting of the samples started at below −5 °C, which was at least 15 °C lower than the starting temperature of melting of normal human meibum. Also, the first major peak of the thermogram had two shoulders at T = 8 °C and 12 °C.

*Subject #2.* Ocular features of the abnormal subject #2 and the preliminary findings on the subject’s abnormal meibum were described in our recent pilot paper^44^. Here, we present results of a more detailed analysis of chemical and biophysical properties of the subject’s MG secretion. Superficially, meibum of Subject #2 looked similar to normal meibum shown in Figures 9A and 9B. However, the sample differed from the normal human meibum in the relative balance of saturated and unsaturated WE, which was shifted toward the former, and in the WE/(Chl + CE) ratio (Figure 9B), which was lower in the abnormal meibum (Figure 9C). The mass spectra of the three major groups of saturated WE (SWE), and MUWE and DUWE are shown in Figure 9D and 9E. Once the LC/MS data for individual WE were integrated, a repeating pattern emerged – the abnormal meibum had a considerably higher proportion of SWE, and lower proportion of MUWE and DUWE regardless of the WE carbon chain length (Figure 9F).

**Figure 9.**
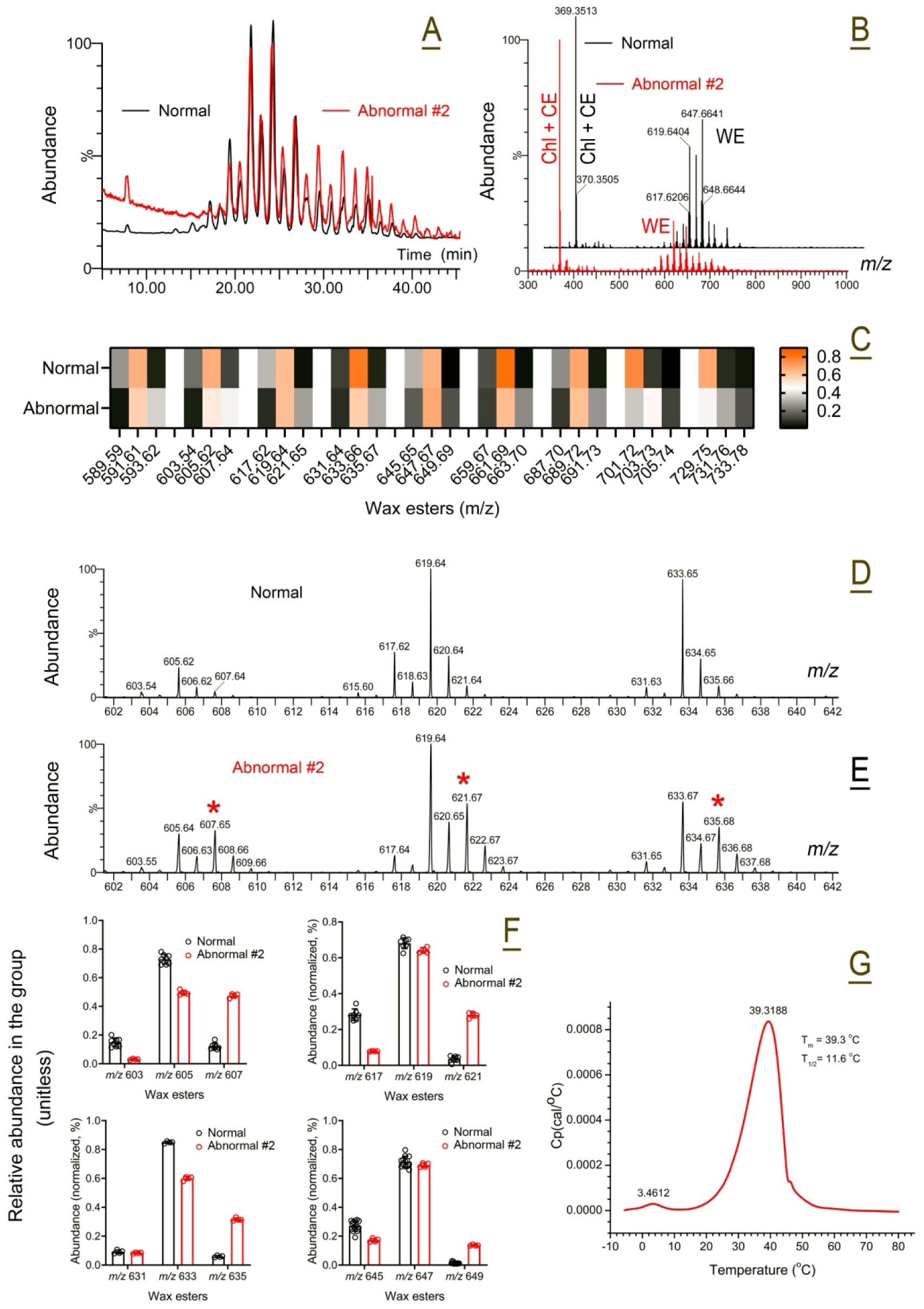
Compositional and functional characterization of abnormal human meibum #2. *<u>Panel A.</u>* Total APCI-PIM ion chromatograms of representative abnormal #2 (orange trace) and normal (black trace) meibum samples obtained as shown in Figure 1, Panel A. Note high similarities in the elution profiles of the samples. Results of an initial LC/MS characterization of the abnormal meibum #2 was partially presented in our recent report^44^. *<u>Panel B.</u>* Observation APCI-PIM mass spectra of abnormal human sample #2 (in orange) and normal meibum (in black). Note an apparently higher (Chl +CE)/WE ratio in the abnormal sample compared with the normal one. *<u>Panel C.</u>* A heatmap of major WE in meibum of normal and abnormal meibum. The compounds are grouped according to their carbon chain lengths. The trios of the compounds are arranged in the following order: diunsaturated, monounsaturated, and saturated WE. The compounds are normalized in each group. The total amounts of three compounds in each group equals 1. Their relative ratios are color-coded: the lowest amounts are shown in black, while the highest – in orange. *<u>Panel D.</u>* A fragment of an APCI-PIM mass spectrum of normal human sample that shows three major monounsaturated wax esters with *m/z* 605, 619, and 633. *<u>Panel E.</u>* A fragment of an APCI-PIM mass spectrum of abnormal human sample #2 that shows upregulation of saturated wax esters with *m/z* 607, 621, and 635. *<u>Panel F.</u>* Extracted ion chromatograms (EIC) of saturated, mono-, and di-unsaturated wax esters were obtained as shown in Figure 1, Panel D, and then integrated to determine the relative balance of the lipids. The data demonstrated a decline in diunsaturated wax esters and upregulation of the saturated compounds, making meibum more solid due to higher melting temperatures of the latter. *<u>Panel G.</u>* Abnormal meibum #2 had a T_m_ of about 39 °C, which is 10 degrees higher than the norm, but similar T_1/2_ value of about 12 °C.

Cumulatively, these changes resulted in a considerable shift in the melting curve of abnormal meibum to higher temperatures with the main T_m_ of 40 ± 1 °C, but virtually the same T_1/2_ of 11.5 ± 1 °C (Figure 9G). Unlike abnormal meibum from Subject **<u>1</u>**, no visible shoulders or peak separation was observed in Subject **<u>2’s</u>** thermograms except for a minor feature at T_m_ of ∼3.5 °C – the secretion remained cohesive regardless of the temperature.

### 2.6. Deconvolution of thermograms

Experimentally obtained thermograms were numerically deconvoluted as described earlier for human meibum samples and WE model mixtures^37^. Deconvolution of the thermotropic peaks can be a useful tool for detecting and visualizing distinctive lipid domains within study samples^45, 46^. Depending on the complexity of the thermograms, the routine allowed for a reasonably accurate approximation of the meibum melting curves using multi-component models with 3 to 8 calorimetric/lipid domains (Figure 10), though calculation of true ΔH_v_ and ΔH values was impossible because of the current lack of data on average molecular weights of animal meibum. Rather complex melting behavior of meibum produced by *Awat2^−/−^, Soat1^−/−^,* and *DKO* mice and abnormal human subject 1 with several clearly visible transition peaks, and very broad (in comparison with pure lipids) melting peaks of other types of samples, can indicate a multistep ***Crystal*** *(solid)∀* ***Liquid crystal*** *(several forms; partially melted)∀* ***Liquid*** *(fully disorganized)* transitions in the samples upon heating, or existence of separate poorly mixing lipid domains (rafts, pools, microcrystals, etc) enriched with different types of lipids that melt at distinctively different temperatures and independently from each other. Establishing the nature of these transitions and/or lipid aggregates goes beyond the scope of this paper and should be addressed in future studies. However, the changes in the apparent melting characteristics of meibum induced by mutations, differences between studied species, and normal and abnormal human meibum, are discussed below.

**Figure 10.**
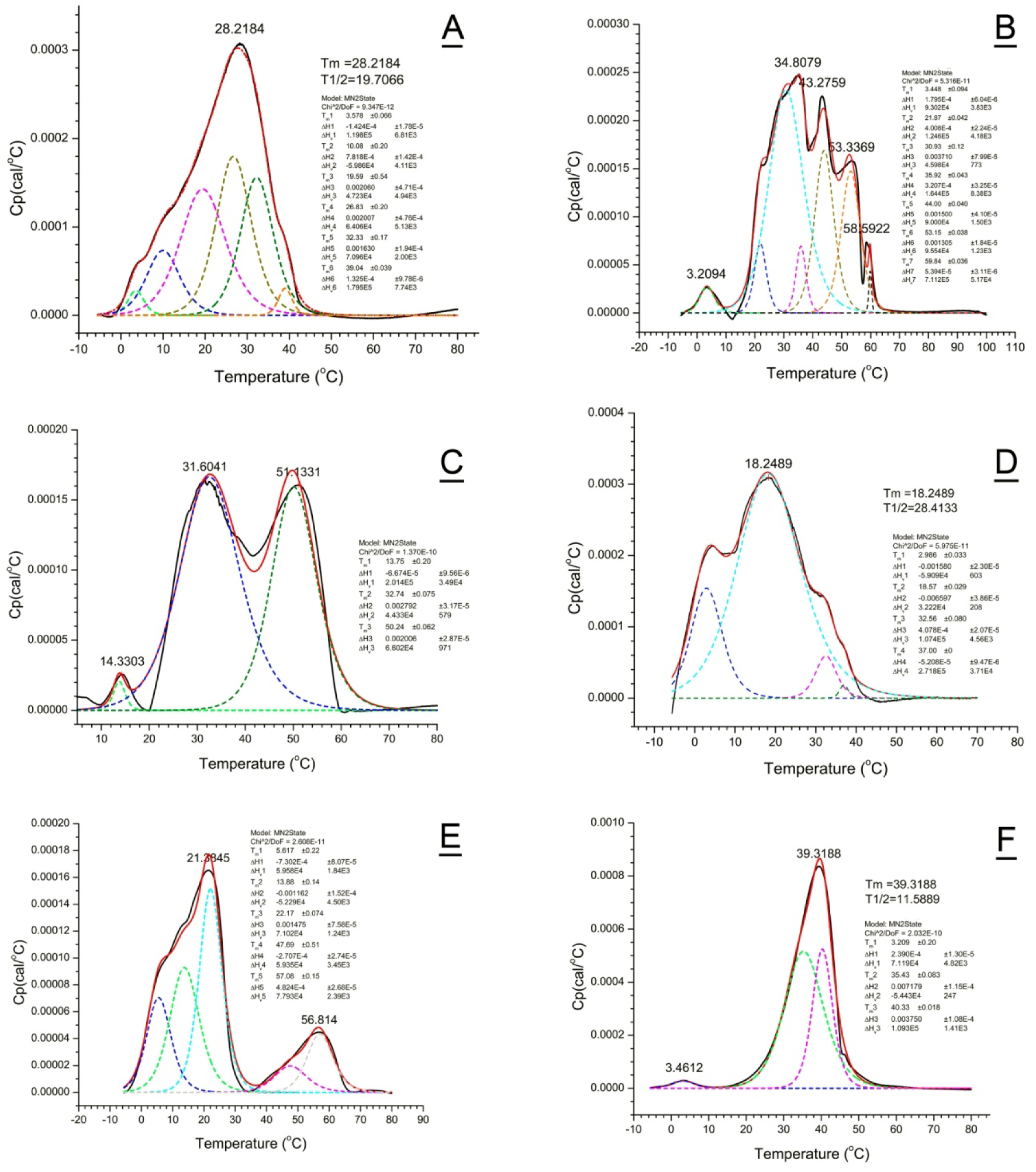
Deconvolution of thermograms reveals multiple phase transitions/calorimetric domains in the samples of wild type and mutant mice and abnormal human meibum #1 and #2. *<u>Panel A.</u>* Wild type meibum. *<u>Panel B.</u> Awat2^−/−^*meibum. *<u>Panel C.</u> Soat1^−/−^*meibum. *<u>Panel D.</u> Sdr16c5^−/−^/Sdr16c6^−/−^ DKO* meibum. *<u>Panel E.</u>* Abnormal human meibum #1. *<u>Panel F.</u>* Abnormal human meibum #2. Black trace – experimental thermogram; red trace – computed thermogram; broken colored lines – deconvoluted individual calorimetric peaks computed using the MN2State model.

#### 2.6.1. WT mouse meibum

A rather wide, but cohesive, melting peak of *WT* mouse meibum with an average T_m_ of 28.2 °C and several shoulders(Figure 10A) implied either a fairly good miscibility of mouse Meibomian lipids that melted uniformly as whole, or the existence of several calorimetric domains that had close T_m_’s. To approximate the shape of *WT* thermograms, up to six T_m_ were needed with three major transitions, rounded to first decimals, at 19.6 ± 0.5, 26.8 ± 0.2, and 32.3 ± 0.2 °C.

#### 2.6.2. Awat2^−/−^ mouse meibum

Meibum of *Awat2^−/−^* mice produced the most complex thermograms of all tested animals being highly incoherent (Figure 10B), which required at least seven different T_m_’s 3.5 ± 0.1, 21.9 ± 0.1, 30.9 ± 0.1 (major), 35.9 ± 0.1, 44.0 ± 0.1 (major), 53.1 ± 0.1 (major), and 59.8 ± 0.1 °C. The temperature at which *Awat2^−/−^* meibum completely melted exceeded 65 °C.

#### 2.6.3. Soat1^−/−^ meibum

Deconvolution of thermograms of *Soat1^−/−^* meibum produced three transition peaks with T_m1_— T_m3_ of 13.8 ± 0.2 (minor), 32.74 ± 0.1 °C (major), and 50.2 ± 0.1 °C (major), correspondingly (Figure 10C). Interestingly, the first T_m1_ was close to the shoulder visible in the thermograms of *WT* mice, while T_m2_ was the close to the main T_m_ of the latter and that of normal human meibum. The most obvious differentiating feature was the third main transition peak with an extremely high T_m3_ of ∼50-51 °C, which is likely to be caused by large amounts of free Chl in the abnormal meibum. The *Soat1^−/−^* meibum became completely liquefied only at temperatures above 60 °C.

#### 2.6.4. DKO meibum

The experimental thermotropic peaks of *DKO* meibum were adequately deconvoluted to produce 4 main superimposed transitions with T_m1_ to T_m4_ of ∼3.0, 18.6, 32.6, and 37.0 °C (Figure 10D). The samples were completely melted by ∼41 ± 2°C.

#### 2.6.5. Canine meibum

Deconvolution of the canine thermograms (not shown) resulted in three major transition peaks T_m1_ = 3.4 ± 0.4 °C, T_m2_ = 23.6 ± 0.1 °C, T_m3_ = 29.1 ± 0.1 °C. Thus, close lipidomic profiles of canine and human meibum correlated well with their close melting characteristics (this paper and ^37^).

#### 2.6.6. Human meibum

Deconvolution of the thermograms of normal human meibum was achieved with three different T_m_^37^, while abnormal meibum #1 required seven (Figure 10E). Numeric deconvolution of the thermogram produced five major components with T_m_ of 5.6 ± 0.2, 13.9 ± 0.2, 22.2 ± 0.1, 47.7 ± 0.5, and 57.1 ± 0.2 °C (all major peaks). The first experimental transition peak with an apparent T_m_ of 21.4 °C could be described well as a superimposition of two peaks with T_m_ values of 13.9 and 22.2 °C. The second largest peak with T_m_ of ∼56.8 °C was also a combination of at least 2 peaks with T_m1_ of 47.7 °C and T_m2_ of 57.1 °C. The nature of these thermograms will be discussed later in the manuscript.

Numerical deconvolution of the thermograms of abnormal meibum #2 resulted in three T_m_ of 3.2 ± 0.2 (minor), and two major peaks at 35.4 ± 0.1 and 40.3 ± 0.1 °C (Figure 10F). Generally, the only difference between this sample and normal human meibum was a considerable increase in the major T_m_ of the abnormal sample, though the overall shape of the peak and its T_1/2_ remained quite similar. Establishing the mechanisms of these reversible rearrangements of lipid aggregates upon heating of normal and abnormal meibum goes beyond the scope of this manuscript and should be attempted in future experiments.

### 2.7. Transcriptomic analysis

Comparative transcriptomic analysis of mouse MG and corneas revealed that most of the genes of interest were barely expressed in the latter. Specifically, the gene expression levels (all shown as the MG/cornea ratios on a log 2 scale) were as follows: *Awat2* (16.8/4.6; *p* < 10^−16^; FDR < 10^−12^), *Elovl3* (17.0/4.5; *p* < 10^−15^, FDR < 10^−11^), *Soat1* (17.3/7.5; *p* < 10^−8^; FDR <10^−6^), *Sdr16c5* (13.1/4.6; *p* < 10^−14^; FDR < 10^−10^), and *Sdr16c6* (16.8/4.4; *p* < 10^−14^; FDR < 10^−10^), while a panel of house-keeping and reference genes^47^ – *Chmp2A* (11.6/12.0), *Emc7* (12.2/10.9), *Gpi1* (13.6/12.4), *Psmb4* (8.6/8.1), *Rab7* (13.8/14.0), *Reep5* (13.0/13.5), *Vcp* (14.7/13.6), and *Vps29* (8.2/8.1) – showed much smaller differences.

## 3. DISCUSSION

Thermotropic properties of meibum determine whether it can fulfill its physiological roles such as being a protector of the eye from the environment, among other functions. To protect the eye, meibum, first, needs to be expressed from the orifices of the exocrine MG (which is mostly a passive process, possibly facilitated by blinking), and, second, spread across the ocular surface and form the protective tear film lipid layer (TFLL; Figure 11). This two-step process critically depends on several factors, among which the eyelid temperature, the ocular surface temperature (OST), and the melting characteristics of meibum, which determine its expressibility from MG’s ducts and ductules, will be discussed below.

**Figure 11.**
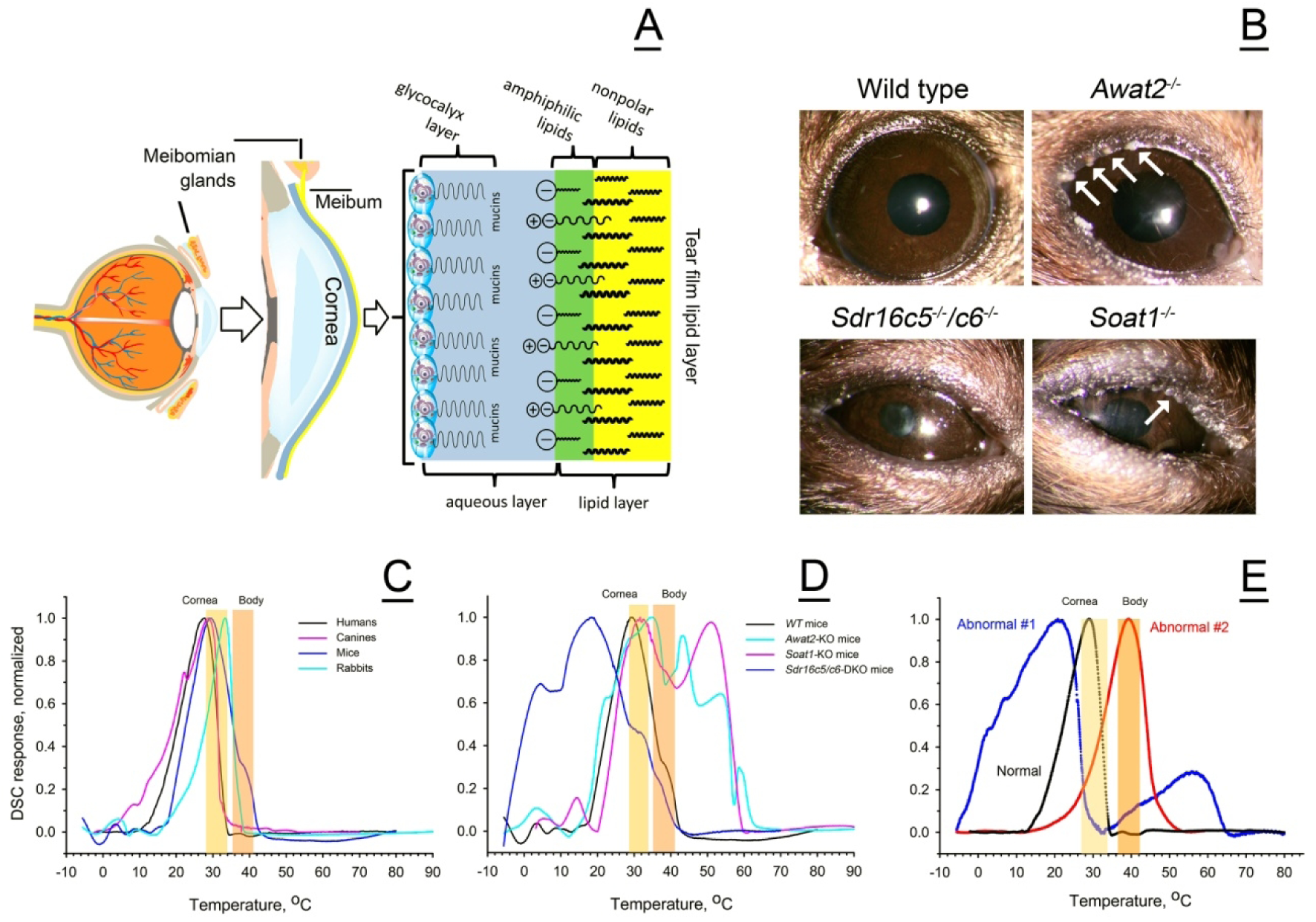
Physiological role of meibum and the impact of its chemical composition on the ocular phenotype. *<u>Panel A.</u>* The eye, cornea, Meibomian glands, and the multilayered/multi-component tear film – a schematic presentation. Meibomian lipids form the most hydrophobic, outermost sublayer of the tear film. Adopted and modified from^35^. *<u>Panel B.</u>* Ocular phenotype of *WT* and mutant mice. Note accumulation of lipid debris around the eye opening, labeled with white arrows. *<u>Panel C.</u>* Comparison of the thermotropic peaks of normal human and animal meibum. Typical cornea temperature ranges are highlighted in yellow, while typical eyelid/body temperature – in pink. *<u>Panel D.</u>* Comparison of wild type and mutant mouse meibum. Note that a large portion of *Soat1^−/−^* and *Awat2^−/−^* meibum does not melt at physiologically normal ocular surface and body temperatures, while *Sdr16c5^−/−^/Sdr16c6 ^−/−^ DKO* meibum starts to melt at below 0 °C. *<u>Panel E.</u>* Thermograms of three types of human meibum samples – normal (black trace), low-temperature-melting type abnormal #2 (blue trace), and high-temperature-melting abnormal #1 (red trace).

Under normal conditions, the human eyelid temperature is between 33 °C and 37 °C^48–50^, and can be somewhat lower or higher, depending on the subject’s health status and environmental conditions. The OST in various corneal regions were reported to be between 30 and 31 °C^51^, 34.3 ± 0.7 °C^38^, 35-35.5 °C^52^, and 37.1 ± 0.1 °C^53^, with a tendency to decline due to the tear film evaporation^54–56^. Thus, prolonged periods of non-blinking ^38, 52, 54^ and high and low air temperatures^57^, among other factors that cause cooling of the ocular surface and adnexa below physiological limits, may facilitate meibum solidification on the ocular surface and/or in the MG ducts, and hinder its expressibility from MG. Higher body and OST, in contrast, would make meibum more liquid than normal. Similarly, a drop in the OST would hamper its spreading across the ocular surface and inhibit formation of the normal TFLL. The in vivo effects of the eyelid temperature on the human meibum expressibility were tested by Nagymihalyi et al^48^, who reported a considerable decline in the delivery of meibum due to low eyelid temperatures, and an increase in meibum delivery/expressibility caused by elevated temperatures. Later, we examined the direct impact of temperature on solidification and melting of normal human meibum using a Langmuir trough and found that Meibomian lipid films become highly elastic upon cooling below the physiological ranges^57^. Our current data on thermotropic properties of human and animal meibum fully support those observations.

However, changes in the eyelid and OST are just two factors that regulate meibum delivery onto the ocular surface: Quality of meibum (or, in the context of this paper, its melting characteristics) is another factor that is critically important for the ocular surface health. In case of abnormal mouse meibum, the general ocular surface phenotypes of *Awat2, Soat1, Elovl3,* and *DKO* mice are highly abnormal (Figure 11 and ^8–13, 58^). The visible changes in the eye ellipticity, signs of inflammation of the eye and adnexa, neovascularization of the cornea, accumulation of lipid deposits on and around the eye, abnormal tearing, and other pathologies were observed in the knockout mice, and replicated those in human patients with MGD and/or DE. Importantly, these genes, being major genes of mouse and human MG, are not expressed in the cornea (Section 2.7 and Figure 11), thus making a direct impact of abnormal meibum on the ocular surface homeostasis due to its altered physical properties and biochemical composition a highly plausible mechanism. As DSC experiments demonstrated (Figures 5D, 11F,Table 2, and^37^), normal human meibum is partially melted at the normal corneal temperature of around 30 °C to 34 °C, and almost completely – at the normal body temperature of 36 °C to 37 °C. In these conditions, meibum is easily expressible from the MG orifices and freely flows across the ocular surface. On the other hand, meibum with abnormally high T_m_ (such as that from human subject #2), will not be easily expressible from the MG orifices, and will stagnate in the MG acini and ducts, lessening the protection of the eye. Moreover, even partially melted meibum with a large pool of unmelted components (as shown in Figure 11E) will not spread evenly across the ocular surface, and will form irregular patches/droplets of lipids seen, e.g., in the tear film of abnormal patient #1^43^. Thus, the bimodal nature of thermograms of abnormal subject #1 may explain the subject’s normal tear film breakup times, due to the components that melt at ∼21 °C, but unusual morphology of the TFLL with large lipid rafts that are clearly visible upon slit lamp examination and do not melt at normal physiological corneal temperatures.

Close similarities between human, canine and *WT* mouse meibum were expected to result in close thermotropic properties of the three secretions, with, possibly, slightly lower T_m_ of the mouse meibum due to slightly shorter carbon chain lengths of the major mouse wax esters (such as C_44_H_86_O_2_ in humans and C_42_H_82_O_2_ in mice^7^). However, the apparent T_m_ of mouse meibum, 28 ± 1 °C, is virtually the same as the human one, but its T_1/2_ is a somewhat broader 19 ± 2 °C (Figure 2 and Table 1). The broadening of the T_1/2_ was caused mainly by the ascending part of the melting curve, which showed a first sign of a phase transition at around 0 °C. Interestingly, the upward section of the mouse thermograms showed a few shoulders which were either much less pronounced, or non-existent, in human samples, though the downward section of the curve had an almost the same shape and slope as the human one. The most distinctive feature of mouse meibum was a secondary, though minor, transition peak, or a shoulder in some samples, with a T_m_ of around 40 °C. The body temperature of WT mice ranges from 36 to 38 °C^59–61^, with the OST being between 36.5 °C to 36.9 °C^53^. This temperature is in the same range as the human corneal temperature^53^, which may imply that mouse meibum, once deposited onto the ocular surface, may behave the same way as the human one. However, if the actual human corneal temperature is closer to 30 °C to 34 °C^38, 51^, then human meibum deposited onto the ocular surface can become less fluid and more liquid crystal-like.

The canine meibum was found to be close to the human meibum in terms of lipid composition, T_m_ and T_1/2_ (Figure 3). Moreover, the OST in dogs, depending on the experiment and environment, ranged from ∼33 °C to ∼38 °C^62–64^, being the same as human and *WT* mouse OST. Thus, canines could have been considered a suitable model for human meibum studies, if not for the obvious ethical considerations. However, close similarities between human and animal ocular surface physiology and biochemistry can be used for developing proper treatments for veterinarian use, as ophthalmic diseases similar to human ones are a major concern in veterinarian practice^65^.

Inactivating mutations in *Awat2, Soat1, a*nd *Sdr16c5/Sdr16c6* genes caused massive changes in the thermotropic properties of mouse meibum (Figure 11D). Aside from shifting major T_m_ to lower or higher temperatures than the norm, the mutations caused considerable broadening of the meibum melting peaks and/or their disintegration into several clearly visible calorimetric domains. Consistent with visual characteristics of *Soat1^−/−^* and *Awat2^−/−^* meibum as being solid or wax-like at physiological temperatures, DSC demonstrated that both secretions undergo complete melting only at temperatures above 60 °C, which means that at physiological temperatures of 30 °C to 37 °C a large portion of meibum exists in solidified state and cannot effectively interact with the tear film. Furthermore, the bi- and multimodal melting of mutant meibum is indicative of abnormal molecular interactions between Meibomian lipids. The complex modality of melting can be caused either by phase separation within the mixture (i.e. due to the formation of separate domains of different types enriched with specific lipids that melt at different temperatures):

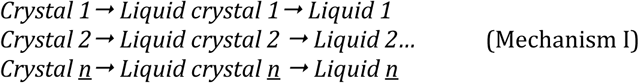

or complex, multistep transitions with formation of several liquid-crystal mesophases:

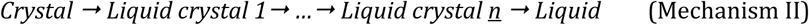

or a combination of both. In normal meibum, these transitions are seamless and produce smooth, cohesive melting curves. Earlier, we demonstrated that the existence of a proper number of lipids of proper types in a mixture is important for generating smooth and cohesive lipid melting curves^37^. It is plausible that the mutation-induced elimination of certain types of lipids from meibum (such as obliteration of all wax esters from *Awat2^−/−^* mice, or cholesteryl esters from *Soat1^−/−^*animals), or abnormal accumulation of certain lipids (such as Chl in the latter, and Chl and short chain and unsaturated wax esters in abnormal human meibum #1) disrupts the normal lipid-lipid interactions in the secretions, causing visible segregation of their calorimetric domains. Though obviously important, establishing the actual mechanisms of these changes necessitates experiments that go beyond the scope of this manuscript and is left for future studies.

Paradoxically, an unexpected result of rather close, but still different, thermotropic properties of human and rabbit Meibomian gland secretions, which are fundamentally dissimilar in terms of their lipid compositions, illustrated that common rules may be applicable to different species: Despite the fact that rabbit meibum has a T_m_ that is ∼ 4 degrees higher than the human one (Figure 4D), their OST is also several degrees higher – about 39.1 ± 0.2 °C^53^. Thus, the melting state and behavior of rabbit meibum on their ocular surface could be similar to the human one, despite being biochemically different.

In summary, inactivation of *Soat1*, *Awat2*, *Sdr16c5/Sdr16c6,* and *Elovl3* genes in mice led to fundamental and complex shifts in their thermograms, which implied selective effects of specific lipids on meibum melting. *Soat1^−/−^* and *Awat2^−/−^* meibum samples had more complex thermograms and higher T_m_ than *WT* meibum, while *Elovl3^−/−^* and *Sdr16c5/Sdr16c6* double knockout meibum displayed an opposite trend. Due to a very low melting point of *Elovl3^−/−^*meibum (T_m_ < 20 °C) and a broad melting curve ^9^, obtaining their DSC was unsuccessful because of the VP-DSC ‘s lowest practical temperature of −5 °C, which was not low enough for recording full melting curves using VP-DSC.

Our data demonstrated that the high complexity of the (bio)chemical composition of normal meibum serves the same goal in all tested species – maintaining its proper physicochemical properties, including, but not limited to, melting characteristics. Major alterations in the balance between main classes of Meibomian lipids, or even changes within individual classes of lipids, may lead to abnormal expressibility, the loss of meibum cohesiveness, and, generally, its ability to protect the eye. DSC seems to be a valuable tool that, together with other approaches, such as LC/MS, can be used in research and clinical practice to detect, characterize, and classify different forms of Meibomian gland dysfunction and other related pathologies, and design strategies for their treatment.

## 4. METHODS

### 4.1. Chemical reagents

The lipid standards (such as saturated and unsaturated wax esters, cholesteryl esters, cholesterol, triacylglycerols, and others) were purchased from Nu-Chek Prep, Inc. (Elysian, MN, USA), Avanti Polar Lipids (Alabaster, AL, USA) and Sigma-Aldrich/MilliporeSigma Chemical Co. (St Louis, MO, USA).

Chromatography/mass spectrometry grade solvents were from Sigma-Aldrich, ThermoFisher (Waltham, MA, USA), or Honeywell-Burdick & Jackson (Muskegon, MI, USA).

### 4.2. Animal and human meibum samples

Animal procedures were approved by the Institutional Animal Care and Use Committee of the UTSouthwestern Medical Center (protocol # 2016-101549, approved on 01/22/2025) and were conducted in accordance with the Association for Research in Vision and Ophthalmology Guidelines. The founder *Sdr16c5^+/−^/Sdr16c6^+/−^*knockout mice on C57BL/6J background were provided by Drs. Natalia Kedishvili and Olga Belyaeva (University of Alabama at Birmingham, Birmingham, Alabama, USA)^13^ and a colony was established at the UTSouthwestern Medical Center (Dallas, TX, USA)^19^. The founder *Soat1*^+/−^ mice on a C57BL/6J background were purchased from the Jackson Laboratory (B6.129S4-*Soat1*^tm1Far/Pgn^; stock #007147; Bar Harbor, ME, USA)^12^. The heterozygous *Elovl3^burf^/GrsrJ* mice were purchased from the same manufacturer (stock no. 024182), and cross-bred with C57BL/6J mice^8^. The *Awat2 ^+/−^*mice on C57BL/6N background (stock number 043484-UCD) were purchased from MMRRC (https://www.mmrrc.ucdavis.edu) and transferred onto C57BL/6J background through multiple generations^10^. All animals used in this study were maintained on a 12-h light/dark cycle and *ad libitum* access to standard commercial laboratory diet (Teklad 2016; Envigo, Somerset, NJ, USA) with free access to water.

All human study sample collection procedures were approved by the Institutional Review Board of the University of Texas Southwestern Medical Center or collaborating institutions. The collection procedures were performed in accordance with the principles of the Declaration of Helsinki. Written informed consents were obtained from all study participants. Human meibum was collected by expressing the secretion from eyelids of healthy, non-DE/non-MGD human volunteers as described before^43^. Meibum of euthanized mice, canines, and rabbits was expressed from the tarsal plates (surgically excised or otherwise) by applying light pressure and collecting the expressed meibum with a metal spatula or a syringe needle with a flat tip, for waxy and solid secretions, or pre-cleaned cellulose tissues (Kimwipes®) for more liquid specimens. If the cellulose wipes were used, the lipids were extracted from the wipes with *n*-hexane/isopropanol (1:1, vol/vol) solvent mixture, the extracts were brought to dryness with a stream of nitrogen. Then, the material was dissolved in iso-propanol (IPA), sealed in glass vials and stored at −80 °C until the analytical experiments. Initially, meibum samples were analyzed using LC/MS, and then compared using the PCA approach. Next, meibum samples were analyzed using DSC, and their T_m_, melting ranges (ΔT), and other parameters were determined as described earlier^37^. In case of complex DSC thermograms, they were numerically deconvoluted, to reveal individual phase transitions/calorimetric domains in the samples. The details of LC/MS and DSC procedures were provided in previous publications, e.g. ^19, 37^, and will be briefly explained in the following sections.

### 4.3. Liquid chromatography – mass spectrometry

Meibum samples were analyzed using reverse phase LC/MS using an M-Class UPLC/Synapt G2-Si system equipped with a C8 or C18 BEH Acquity chromatographic columns (all from Waters Corp., Milford, MA, USA) as described earlier^19^ and shown in Figure 6. The elution of the analytes was conducted using a ternary gradient of IPA, acetonitrile, and 10 mM aqueous ammonium formate with a C18 BEH column, or an isocratic elution with a C8 BEH column. The lipids were identified on the basis of their *m/z* values, fragmentation patterns in MS/MS or MS^E^ mode, LC retention times, and comparison with authentic chemical standards, such as WE, Chl, CE, TAG, and other lipids, if available. The analytes were detected either APCI, or electrospray ionization modes^7, 19^. The raw data were acquired in the *m/z* 100 to *m/z* 2,000 range with 5- to 10 mDa accuracy, and individual lipids were analyzed as EIC. After their integration, relative abundances of the individual analytes were compared as fractions of the total lipid peaks, or as fold changes in meibum samples of different types.

Then, the data was exported to the Progenesis QI and EZinfo software packages (also from Waters) for final analyses using an unbiased, untargeted multivariate PCA and/or PLS-DA approaches.

### 4.4. Differential scanning microcalorimetry

The meibum samples dissolved in IPA were loaded in the sample cell of the differential scanning microcalorimeter (VP-DSC from MicroCal/GE, Piscataway, NJ, USA) using a microsyringe and the solvent was evaporated under a stream of dry ultrapure nitrogen, while the reference cell was left empty. Typically, between 100 and 300 μg of dry lipid material were loaded for each DSC experiment. Knowing the exact amount of loaded material was not essential for this study as the results were compared after normalization of DSC thermograms of all samples (the DSC response from 0% to a maximum of 100%). However, larger amounts of the samples radically improved the signal-to-noise ratio and provided more reproducible results. Then, the cells were sealed, and the heating-cooling cycles from −9 °C to +80 °C were started at a typical scan rate of 2 °C/min. Fifteen consecutive thermograms obtained for a representative canine sample are shown in Figure 12A as overlaid curves, 14 of which (from #2 to #15) demonstrated high reproducibility of the experiments. The first heating/cooling cycle is considered to be a preparatory one^37^ and has been excluded from further analyses. For actual meibum melting experiments, heating-cooling cycles were repeated 5 times or more. Then, the scans from #2 and up were averaged using MicroCal/Origin 7 software supplied with the VP-DSC instrument, the baseline correction was performed using the baseline correction algorithm of the software, and the parameters of the melting curves, such as **T_m_**, peak areas **A**, and **T_1/2_,** were determined (Figure 12B). These parameters were calculated directly from the thermograms using the “Pick Peaks” and “Integrate from zero” commands of the Origin/MicroCal software. Other approaches and/or programs can be used for the same purposes^66, 67^. Where applicable, the thermograms were deconvoluted using “MN2State: Cursor Initiated” routine of the Origin/MicroCal software, and the T_m_ values of the deconvoluted melting peaks were determined. Also calculated were *apparent* van’t Hoff enthalpy changes (**ΔH_V_**) and calorimetric transition enthalpies (**ΔH**). The use of apparent ΔH_V_ and ΔH was dictated by uncertainty in the estimation of the average molecular weights of all but human study samples and are provided here only to illustrate the ability of the tested approach to simulate the actual thermograms of meibum.

**Figure 12.**
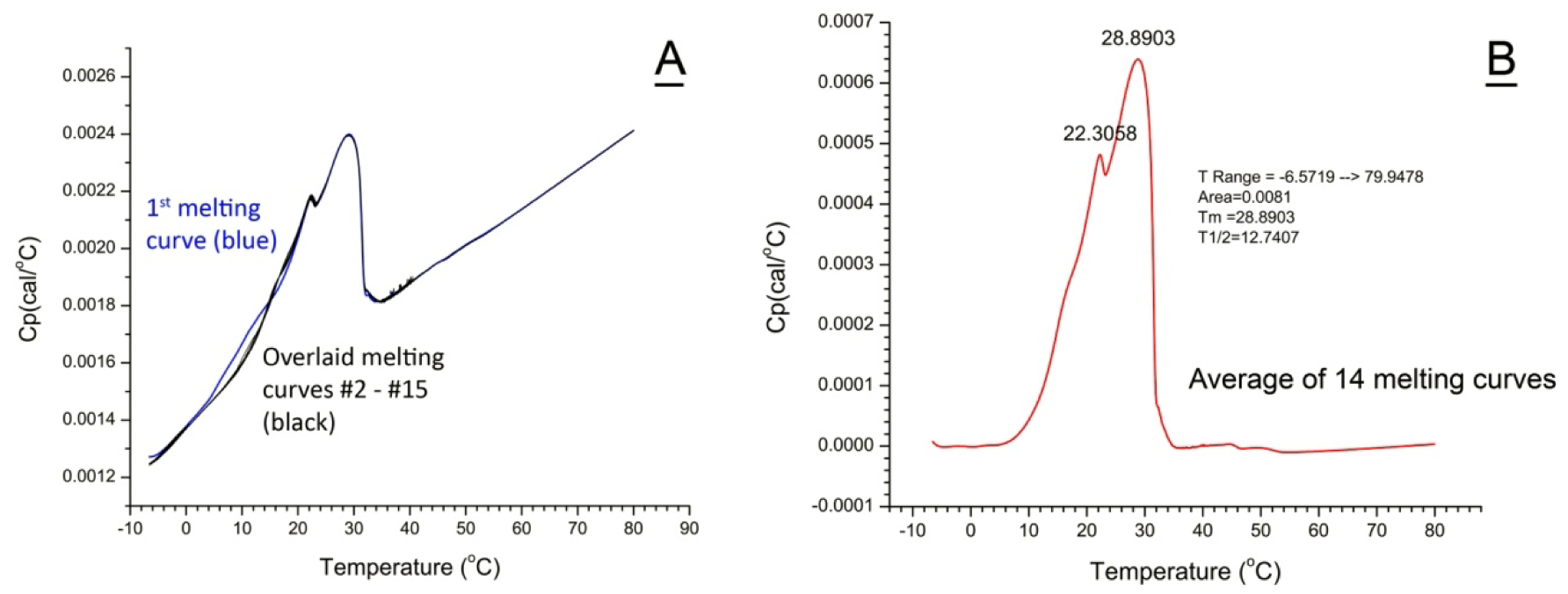
Differential scanning microcalorimetric analysis of normal canine meibum demonstrated high reproducibility of the results and stability of the sample during multiple heating-cooling cycles. *<u>Panel A.</u>* Fifteen sequential scans of the same meibum sample demonstrate high reproducibility of the analyses. Scan #1 (in blue) is considered a preparatory step. Raw unprocessed scans are shown. *<u>Panel B.</u>* A thermogram obtained after averaging scans 2 to 15 from Panel A and correcting for the baseline. The peak area, transition temperature T_m1_ and T_m2_, and T_1/2_ width of the peak were calculated as described in the Methods section.

### 4.5. Transcriptomic analysis of mouse tarsal plates and corneas

The expression levels of *Awat2*, *Soat1*, *Elovl3,* and *Sdr16c5/Sdr16c6* genes in mouse corneas and MG were determined at the UTSW Microarray and Immune Phenotyping Core using Clariom D microarrays (from Affymetrix, Santa Clara, CA, USA), and the data were exported to the Expression Console and Transcriptome Analysis Console (TAC, from the same manufacturer)for quantitation as described earlier^58^. An empirical Bayes ANOVA (eBayes) algorithm was used for ranking differentially expressed genes. Based on industry-standard settings, the linear fold changes in gene expression patterns that satisfied the following criteria— (−2) ≥ LFC ≥ (+2) and *p* ≤ .05—were considered to be statistically significant.

## AUTHORSHIP CONTRIBUTION STATEMENT

**I.A.Butovich**: Conceptualization; experimentation; method development; data acquisition and analysis; supervision the project; project administration; funding acquisition; writing and reviewing the manuscript

**J.C.Wojtowicz**: Clinical testing of human subjects and human sample collection; experimentation; data analysis; reviewing the manuscript

**A. Wilkerson**: Animal husbandry and care; animal sample collection

**S. Yuksel**: experimentation; method development; Animal husbandry and care; animal sample collection; data analysis.

## DATA AVAILABILITY

All data used in the manuscript are included as graphs, tables, and Supplemental Materials. For additional inquires, please contact the corresponding author (IAB) directly.

## ACKNOWLEDGEMENTS

The authors would like to thank Dr. Tomo Suzuki (Kyoto Prefectural University of Medicine, Kyoto, Japan) and Dr. Daniel A. Johnson (University of Texas Health Science Center in San Antonio, San Antonio, TX, USA) for providing abnormal human meibum samples #1 and #2, respectively. A more detailed analyses of these samples is underway and will be reported separately. The results included in this manuscript were partially presented at the ARVO annual conference in Salt Lake City, Utah, in May 2025 (abstract #6077) and in ^44^.

## ABBREVIATIONS

APCI PIM: atmospheric pressure chemical ionization positive ion mode
Chl: free cholesterol
CE: cholesteryl ester(s)
DE: dry eye
DKO: *Sdr16c5^−/−^/Sdr16c6^−/−^* mice
ΔH: calorimetric transition enthalpies
ΔH_v_: van’t Hoff enthalpy changes
DE: dry eye
DiAD: α,ω-diacylated diols
DiHLE: dihydrolanosteryl ester(s)
DSC: differential scanning microcalorimetry
DUWE: diunsaturated wax ester(s)
FA: fatty acid(s)
FFA: free fatty acid(s)
GEE: glyceryl ether ester(s)
LC: liquid chromatography
MS: high resolution time-of-flight mass spectrometry
*m/z*: mass-to-charge ratio (unitless)
MG: Meibomian gland(s)
MGD: Meibomian gland dysfunction
MUWE: monounsaturated wax ester(s)
OST: ocular surface temperature
PCA: Principal Component Analysis
PLS-DA: Partial Least Squares Discriminant Analysis
SE: steryl ester(s) (non-cholesterol-based)
SG: sebaceous gland(s)
SWE: saturated wax ester(s)
T_1/2_: the width of the transition at half-height of the melting peak
TAG: triacylglycerol(s)
TF: tear film
TFLL: tear film lipid layer
T_m_: transition temperature
WE: wax ester(s)
WT: wild type

